# NIR-II squeezed light-field microscopy enables high-speed volumetric imaging of deep-tissue dynamics *in vivo*

**DOI:** 10.64898/2026.08.13.744709

**Authors:** Do Young Kim, Zihan Zang, Eric Y. Lin, Ruixuan Zhao, Jing Wang, Chen Yuan Kam, Tzung K. Hsiai, Ellen M. Sletten, Liang Gao

**Affiliations:** Department of Bioengineering, Henry Samueli School of Engineering and Applied Science, University of California, Los Angeles, Los Angeles, CA, USA; Department of Chemistry and Biochemistry, University of California, Los Angeles, Los Angeles, CA, USA; Division of Cardiology, Department of Medicine, David Geffen School of Medicine, University of California, Los Angeles, Los Angeles, CA, USA; Division of Dermatology, Department of Medicine, David Geffen School of Medicine, University of California, Los Angeles, Los Angeles, CA, USA; Department of Molecular, Cell, and Developmental Biology, University of California, Los Angeles, Los Angeles, CA, USA; Eli and Edythe Broad Center for Regenerative Medicine and Stem Cell Biology, University of California, Los Angeles, Los Angeles, CA, USA; Greater Los Angeles VA Medical Center, Los Angeles, CA, USA

## Abstract

High-speed three-dimensional imaging in scattering biological tissues remains challenging because volumetric microscopy generally requires scanning, whereas snapshot light-field approaches divide limited detector pixels among multiple views. This constraint is particularly severe in the second near-infrared window (NIR-II), where InGaAs cameras have small sensor formats and high detector noise. Here we introduce NIR-II squeezed light-field microscopy (NIR-II SLIM), which optically rotates and compresses multiple perspective views before detection, allowing efficient use of camera pixels while retaining complementary spatial information for three-dimensional reconstruction. NIR-II SLIM acquires up to 600 volumes s⁻¹ with a reconstructed lateral sampling grid of 512 × 512 pixels. We use this method for label-free imaging of cardiac dynamics in pigmented late-larval zebrafish, resolving chamber deformation and millisecond-scale atrioventricular-valve motion, and for NIR-II fluorescence imaging of vascular and lymphatic transport in mice. NIR-II SLIM provides a detector-efficient approach for high-speed volumetric imaging of rapid biological dynamics in scattering tissues.

## Introduction

Capturing rapid three-dimensional (3D) biological processes *in vivo* requires imaging modalities that combine high spatial and temporal resolution with sufficient penetration depth. Although fluorescence microscopy has fundamentally transformed the study of biological structure and function, visualizing fast volumetric events deep within scattering tissues remains challenging. Visible light undergoes substantial scattering and absorption in biological tissue, generally restricting high-resolution imaging to superficial regions or requiring invasive surgical procedures or optical-clearing preparations.

These limitations have motivated a shift toward longer optical wavelengths, particularly the second near-infrared window (NIR-II; 1,000–1,700 nm). Reduced tissue scattering in this spectral range can extend imaging depth and improve contrast in intact biological specimens^1–4^. Interest in NIR-II imaging has consequently grown rapidly, accompanied by an expanding repertoire of emitters^5–7^, including single-walled carbon nanotubes^8^, small organic molecules^9^, rare-earth-doped nanoparticles^10^, quantum dots^11^, conjugated polymers^12^, and other inorganic nanomaterials^13^. Together, these probes have enabled important advances in both basic biomedical research and translational applications.

Despite this progress, many widely used *in vivo* NIR-II imaging methods remain limited to two-dimensional (2D) visualization. This constraint is particularly restrictive for intact-organ mechanics, hemodynamics, and lymphatic dynamics, which arise from intrinsically volumetric structures and evolve across both depth and time. In the heart, for example, myocardial wall thickening and torsion, valve-leaflet motion, and intracardiac flow result from coordinated 3D interactions^14,15^. Depth-integrated measurements cannot fully recover three-dimensional chamber geometry or out-of-plane strain, potentially obscuring important features of cardiac development, congenital abnormalities, and drug-induced mechanical dysfunction. Similarly, conventional 2D NIR-II angiography projects vessels at different depths onto a single plane, confounding vascular geometry and complicating measurements of vessel diameter, branching, tortuosity, and depth-resolved perfusion. 2D lymphangiography likewise cannot fully resolve the organization of lymphatic networks or track flow-associated signals through vessels distributed across multiple tissue depths.

Existing approaches to 3D NIR-II imaging generally rely on sequential spatial scanning, as in confocal, spinning-disk confocal^16,17^, and light-sheet microscopy^18,19^. Although effective for static or slowly evolving specimens, scanning inherently constrains volumetric acquisition speed. Even state-of-the-art NIR-II light-sheet systems typically acquire volumes at only approximately 0.4 volumes per second^18^, far too slow to resolve rapid processes such as cardiac motion, vascular hemodynamics, and lymphatic pulsatility. The resulting mismatch between acquisition speed and physiological motion can introduce substantial motion artifacts, distort volumetric reconstructions, and obscure transient events. A fundamentally different imaging strategy is therefore needed to capture fast biological dynamics in intact, scattering tissues with both high speed and high spatial fidelity.

Light-field imaging offers a promising solution by recording multiple angular perspectives simultaneously and computationally reconstructing depth from the disparities among them^20–22^. Because all views are acquired in a single exposure, light-field imaging eliminates mechanical scanning and enables snapshot volumetric imaging. This capability has supported a broad range of biomedical applications in the visible spectrum, including microscopy, endoscopy, and mesoscopy^20^.

Extending light-field imaging to the NIR-II range, however, presents a major detector challenge. Conventional light-field architectures encode both spatial and angular information by dividing the available sensor area among multiple perspective views, creating a direct tradeoff between angular sampling and spatial resolution. This tradeoff is especially restrictive for NIR-II imaging, which typically relies on indium gallium arsenide (InGaAs) cameras. Although InGaAs focal-plane arrays provide sensitivity throughout the NIR-II spectral range, their compound-semiconductor fabrication and hybrid integration with silicon readout circuits increase manufacturing complexity and cost and constrain pixel pitch and array format relative to mature silicon imagers^23,24^. InGaAs cameras also typically exhibit substantially higher readout noise and dark current than silicon-based cameras^25^, which can limit sensitivity under photon-starved NIR-II fluorescence conditions.

These constraints substantially limit the performance of conventional NIR-II light-field microscopes. For example, a recently reported Fourier light-field microscope^26^ used a 640 × 512-pixel InGaAs sensor partitioned into a 3 × 3 array of perspective views. Each view therefore contained only approximately 213 × 170 pixels, substantially reducing the spatial information available for volumetric reconstruction. An effective NIR-II light-field imaging strategy must therefore address two coupled challenges: efficiently encoding multiple angular perspectives on a pixel-limited image sensor and minimizing the noise penalty associated with reading out large numbers of high-noise InGaAs pixels.

To address these challenges, we present NIR-II Squeezed light-field microscopy (NIR-II SLIM), a computational imaging platform that extends the squeezed optical-mapping strategy^27^ recently developed in our laboratory to the NIR-II spectral window. NIR-II SLIM reduces redundant sampling of spatial and angular information, enabling high-resolution volumetric imaging with limited-format InGaAs cameras.

Unlike conventional light-field microscopy, which partitions the detector into a 2D grid of uniformly downsampled views, NIR-II SLIM uses a linear array of Dove prisms and anamorphic relay optics to rotate and compress the perspective images along a single spatial axis. Each subaperture image retains full resolution along its orthogonal, unsqueezed axis. By distributing these high-resolution axes across complementary orientations, the system preserves nearly the same lateral spatial bandwidth as unsqueezed detection while accommodating multiple perspective images within the available InGaAs sensor format.

The squeezed encoding also provides a detector-noise multiplexing advantage analogous to the Fellgett advantage^28^. Under the detector-noise-limited conditions typical of NIR-II fluorescence imaging, conventional light-field microscopy spreads the signal across many InGaAs pixels, whose readout and dark noise accumulate during reconstruction. NIR-II SLIM instead compresses and multiplexes the required spatial and angular information onto fewer detector pixels, reducing the aggregate noise contribution. Thus, squeezing not only accommodates multiple perspective images on a pixel-limited sensor but also improves the signal-to-noise ratio (SNR) under detector-noise-limited conditions.

Combined with the intrinsic snapshot capability of light-field imaging, this efficient encoding enables volumetric acquisition at rates of up to 600 volumes s⁻¹ with a reconstructed lateral sampling grid of 512 × 512 pixels. Compared with state-of-the-art NIR-II light-field imaging methods^26,29^, NIR-II SLIM provides a six-fold increase in volumetric imaging speed and an over four-fold increase in reconstructed lateral pixel count.

We demonstrate these capabilities in biological applications requiring rapid volumetric imaging over reconstructed axial ranges of approximately 0.6 mm in mouse ear vasculature and 3 mm in tumor-draining lymphatics. First, we address a fundamental limitation of zebrafish imaging: beyond approximately 7 days post-fertilization, progressive melanophore pigmentation renders the animals increasingly opaque to visible light. By combining dark-field illumination with the reduced absorption of melanin in the short-wave infrared range, NIR-II SLIM enables label-free 4D cardiac imaging in zebrafish aged 15–25 days post-fertilization. Second, we use micelles of the chromenylium NIR-II fluorophore Chrom7^30^, which exhibits an emission tail extending into the 1,100–1,300 nm detection range, to noninvasively image tumor-draining lymphatic dynamics *in vivo* in mice. NIR-II SLIM tracks bolus transit through mouse ear vasculature at 20 volumes s⁻¹ and resolves lymphatic bolus propagation at 100 volumes s⁻¹ across multiple tissue depths.

Together, these results establish NIR-II SLIM as a powerful platform for high-speed, deep-tissue volumetric microscopy. By combining efficient use of limited detector pixels, improved performance under detector-noise-limited conditions, and snapshot 3D acquisition, NIR-II SLIM enables quantitative visualization of transient physiological processes in their native scattering environments and opens new opportunities for investigating deep-tissue dynamics *in vivo*.

## Results

### Optical configuration

The optical configuration of the NIR-II SLIM system is shown in Fig. 1a. The system can operate in either dark-field or fluorescence imaging mode. In dark-field mode, the sample is trans-illuminated at a large oblique angle, and an imaging objective collects only the light scattered by the specimen. The directly transmitted illumination falls outside of the collection numerical aperture of the objective and is therefore rejected. In fluorescence mode, the sample is illuminated in an epi-configuration, and the emitted fluorescence is collected through the same objective. A dichroic mirror separates the excitation and emission paths.

**Fig. 1.**
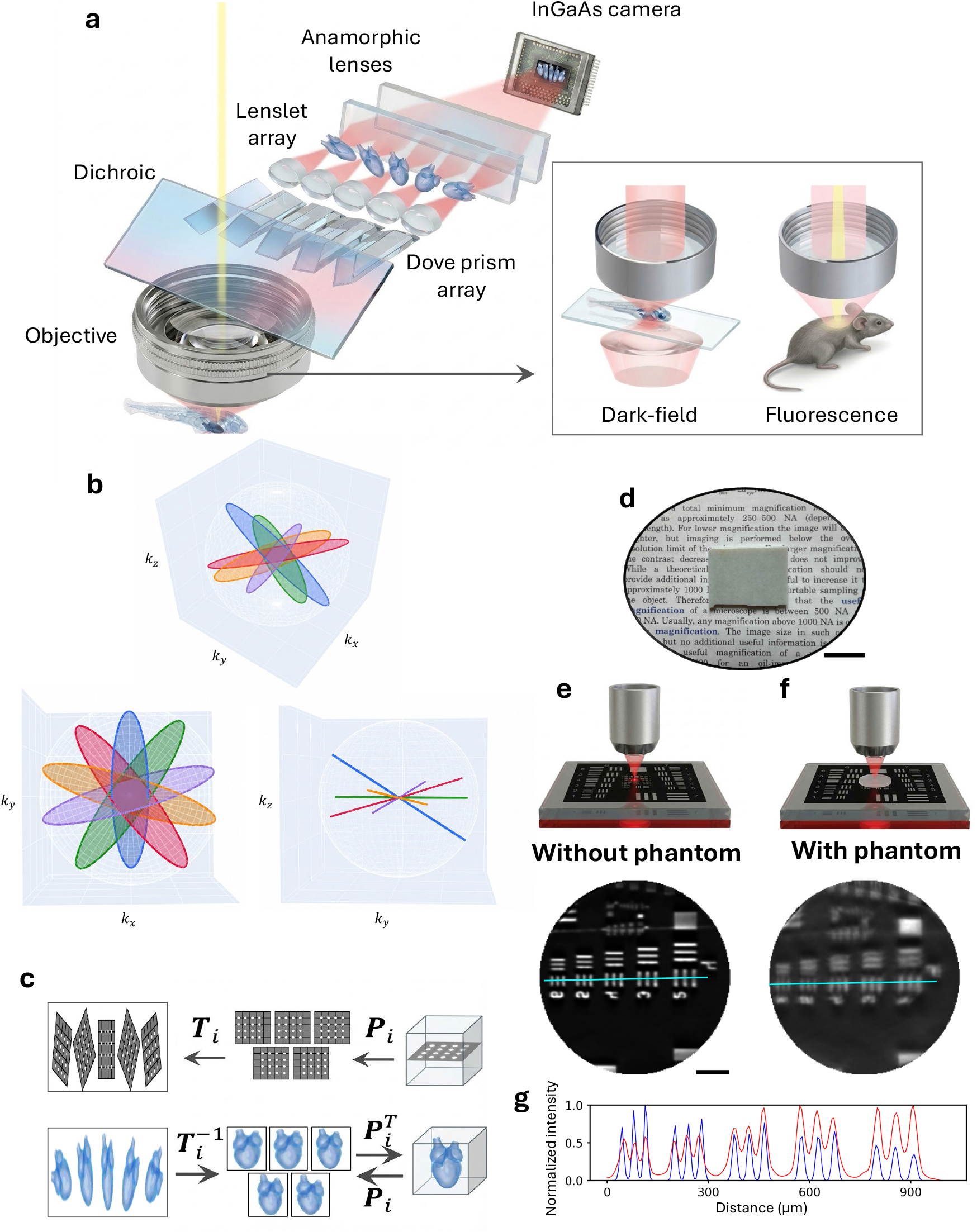
Optical configuration, reconstruction framework, and imaging through scattering tissue phantoms. **a,** Optical configuration of NIR-II squeezed light-field microscopy (SLIM). Light collected by the objective is divided into five subaperture views by a linear Dove-prism and lenslet array, rotated to complementary orientations and anisotropically compressed before detection with an InGaAs camera. The system supports dark-field transillumination for label-free imaging and epi-fluorescence illumination for NIR-II fluorescence imaging. **b,** Fourier-domain representation of the five perspective views; colors denote the five views. Anamorphic compression produces anisotropic spatial-frequency support for each view, while Dove-prism rotation distributes the high-bandwidth axes across complementary orientations in three-dimensional frequency space. A three-dimensional view is shown together with projections onto the *k_x_*–*k_y_* and *k_y_*–*k_z_* planes. **c,** Physics-based forward model and reconstruction framework. Depth-dependent projection operators, *P_i_*, map the three-dimensional object onto the five perspective views, and view-specific affine transformations, *T_i_*, account for rotation, anisotropic magnification, and camera-plane distortions. Reconstruction applies the inverse geometric transformations followed by iterative back-projection using *P_i_* and its adjoint 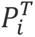. **d,** Photograph of a 2-mm-thick tissue-mimicking optical phantom (QUEL Imaging; μ_a_ = 0.027 mm^-1^, μ_s_′ = 0.42 mm^-1^ at 800 nm) placed over printed text, illustrating its opacity in the visible spectrum. Scale bar, 1 mm. **e, f,** NIR-II SLIM imaging of a USAF resolution target under diffuse 1,300-nm transillumination without (**e**) and with (**f**) the 2-mm-thick phantom. Top, imaging configurations; bottom, computationally refocused images. Cyan lines indicate the positions used for intensity-profile analysis. Scale bar, 200 µm. **g,** Normalized intensity profiles along the lines indicated in **e, f** (blue, without phantom; red, with phantom). Despite reduced contrast and profile broadening through the phantom, Group 4, Element 6 remained resolved, corresponding to approximately 28.5 line pairs mm⁻¹.

On the detection side, the input image is relayed through a linear array of Dove prisms paired with lenslets. Each Dove prism is rotated by a distinct angle about its optical axis, producing a corresponding perspective image rotated by twice the prism angle. The resulting array of angularly distinct perspective images is formed at an intermediate image plane behind the lenslet array.

The perspective-image array is subsequently relayed through an anamorphic optical system composed of two orthogonally oriented cylindrical-lens assemblies. This system applies anisotropic magnification, reducing the image scale by approximately fivefold along one spatial axis while preserving the magnification along the orthogonal axis. The compressed perspective images are then recorded by a 2D InGaAs camera. By squeezing the image array along only one dimension, this optical mapping substantially reduces the detector area and pixel count required to capture the full set of perspective views. A detailed description of the optical design and system configuration is provided in the Methods section (Supplementary Figs. 1 and 2).

The frequency-domain consequence of this squeezed encoding is illustrated in Fig. 1b. Under the Fourier-slice interpretation^31^, each perspective view samples a distinct slice of the 3D spatial-frequency spectrum, with its orientation determined by the corresponding subaperture viewing direction. Anamorphic compression produces anisotropic frequency support, reducing the sampled bandwidth along the squeezed dimension while preserving higher bandwidth along the orthogonal dimension; Dove-prism rotation distributes these preserved high-bandwidth axes across complementary orientations. Consequently, the five subaperture views provide complementary, anisotropically elongated Fourier-domain slices that intersect near the origin but extend along different lateral directions (*k*_x_–*k*_y_ plane). This complementary sampling preserves high spatial-frequency information across multiple orientations while substantially reducing the detector area required for recording the perspective-image array, thereby providing the frequency-domain basis for high-resolution 3D reconstruction from the squeezed measurements (Supplementary Note 1 and Supplementary Figs. 8 and 9).

### Forward model and physics-informed reconstruction

Accurate 3D reconstruction from the compressed measurements requires a forward model that accounts for both the geometric transformations introduced by the optical system and the depth-dependent point-spread functions (PSFs) of the subaperture views. The image-formation process can be decomposed into a depth-dependent light-field projection followed by geometric mapping. However, the Dove prisms and anamorphic relay introduce view-dependent image rotation and anisotropic magnification, breaking the shift invariance typically assumed in conventional light-field microscopy and precluding a single convolutional representation.

To account for these effects, we define an affine transformation ***T****_i_* for each subaperture view *i*. This transformation describes the view-specific rotation, anamorphic magnification, and other geometric distortions between the virtual intermediate image plane and the camera plane. After applying the inverse transformation 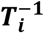, the rectified subaperture image is mapped back to the virtual intermediate image space, where the relationship between the 3D object and each perspective view becomes approximately shift invariant.

We then define ***P****_i_* as the light-field projection operator for the *i*-th subaperture. Each ***P_i_*** maps the 3D sample distribution onto the corresponding perspective image using depth-dependent PSFs measured throughout the imaging volume. The measurement from each subaperture can therefore be expressed as

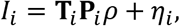

where *I_i_* is the compressed camera measurement associated with the *i*-th perspective view, *ρ* denotes the 3D fluorescence- or scattering-intensity distribution, depending on the imaging modality, and *η_i_* represents measurement noise. The complete forward operator is formed by combining the measurements from all *N* subapertures:

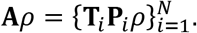

We experimentally calibrate this physics-based forward model by axially scanning a pinhole array through the imaging volume and recording the response of every subaperture at multiple depths (Fig. 1c, top row). These measurements determine the depth-dependent PSFs used to construct **P*_i_***. The positions, orientations, and anisotropic scaling of the pinhole images are simultaneously used to estimate the geometric parameters of **T*_i_***. The calibrated model therefore incorporates the actual optical response of the system, including depth-dependent blur, angular parallax, image rotation, and anamorphic compression.

Reconstruction is formulated as a physics-informed inverse problem in which the recovered volume must remain consistent with the measured data under this calibrated forward model. As illustrated in Fig. 1c (bottom row), the measured subaperture images are first rectified using 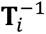 and mapped to the virtual intermediate image space. In this coordinate system, the operators **P***_i_* and their adjoints 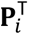 propagate information between the 3D object space and the measured perspective views. At each iteration, the current volume estimate is projected through the experimentally calibrated imaging model, compared with the measurements, and updated by back-projecting the resulting discrepancy. The reconstruction therefore explicitly enforces the physical processes governing image formation rather than relying solely on generic image priors or statistical correlations.

### Imaging through scattering tissue phantoms

To evaluate the ability of NIR-II SLIM to preserve spatial resolution through strongly scattering media, we imaged a resolution target through a tissue-mimicking optical phantom. A 2-mm-thick tissue phantom (QUEL Imaging; *μ_a_* = 0.027 mm^−1^, *μ_s_*′ = 0.42 mm^−1^, specified at 800 nm) was placed directly above a USAF resolution target. As shown in Fig. 1d, the phantom is highly opaque in the visible spectrum, substantially obscuring the printed text beneath it. For NIR-II imaging, the resolution target was trans-illuminated with diffuse light centered at 1,300 nm and imaged through the phantom using NIR-II SLIM.

We first imaged the resolution target without the phantom to establish a reference measurement (Fig. 1e). We then repeated the measurement after placing the 2-mm-thick phantom over the target using the same imaging configuration. Computational refocusing of the reconstructed SLIM volume recovered the target features despite the intervening scattering layer (Fig. 1f).

A quantitative comparison of the intensity profiles acquired with and without the phantom is shown in Fig. 1g. Although the phantom reduced image contrast and broadened the intensity profiles, the periodic structure of the resolution target remained clearly distinguishable. Notably, NIR-II SLIM resolved Group 4, Element 6 bars through the 2-mm-thick phantom, corresponding to a spatial frequency of approximately 28.5 line pairs mm⁻¹. Calibration-pinhole axial responses and localization errors, center-depth lateral resolution, and the depth- and field-dependent variation of measured pinhole-image widths are characterized in Supplementary Figs. 3, 4, and 5, respectively. These results demonstrate that NIR-II SLIM preserves fine spatial detail through millimeter-thick scattering media, supporting its potential for high-resolution volumetric imaging in scattering biological tissues.

### Label-free 4D cardiac imaging in late-larval zebrafish

Although zebrafish provide a powerful vertebrate model for investigating cardiac development and disease^32–35^, cardiac imaging becomes increasingly challenging during postembryonic growth. Conventional optical studies have therefore focused predominantly on embryos and early larvae, when the heart remains readily accessible to visible-light microscopy^36,37^. At later stages, progressive melanophore pigmentation increasingly attenuates visible light and obscures internal cardiovascular structures. At the same time, continued cardiac growth, rotation, and chamber remodeling increase the 3D extent of the heart, making single-plane measurements increasingly sensitive to specimen orientation and focal-plane selection. Late-larval zebrafish therefore occupy an important imaging gap: they are no longer sufficiently transparent for conventional high-speed visible-light microscopy yet remain too small for modalities such as ultrasound or MRI to provide comparable spatial resolution^38^.

This limitation is particularly relevant between approximately 15 and 25 days post-fertilization (dpf), when substantial postembryonic cardiac maturation occurs. During this period, atrioventricular (AV) valve leaflets continue to elongate and subsequently undergo extracellular matrix deposition and thickening^39^. Coronary and cardiac lymphatic vessels likewise begin to form only at juvenile stages, from approximately 1–2 months post-fertilization^32,33^, so that valvulogenesis, coronary angiogenesis, and cardiac lymphangiogenesis all unfold within a window that is inaccessible to visible-light microscopy yet below the resolution of ultrasound or MRI. Consequently, important developmental processes and progressive cardiac phenotypes that emerge beyond the embryonic stage remain difficult to interrogate dynamically *in vivo*. Because melanin absorption is reduced at longer wavelengths, NIR-II imaging provides a means to access these increasingly pigmented animals. We therefore used NIR-II SLIM to perform label-free volumetric imaging of cardiac dynamics in 20-dpf wild-type and weak atrium (*wea*; *myh6*^−/−^) zebrafish (Fig. 2). In *wea* mutants, loss of atrial myosin heavy-chain function disrupts atrial myofibrillar organization and contractility, while ventricular contraction remains comparatively preserved^40,41^.

**Fig. 2.**
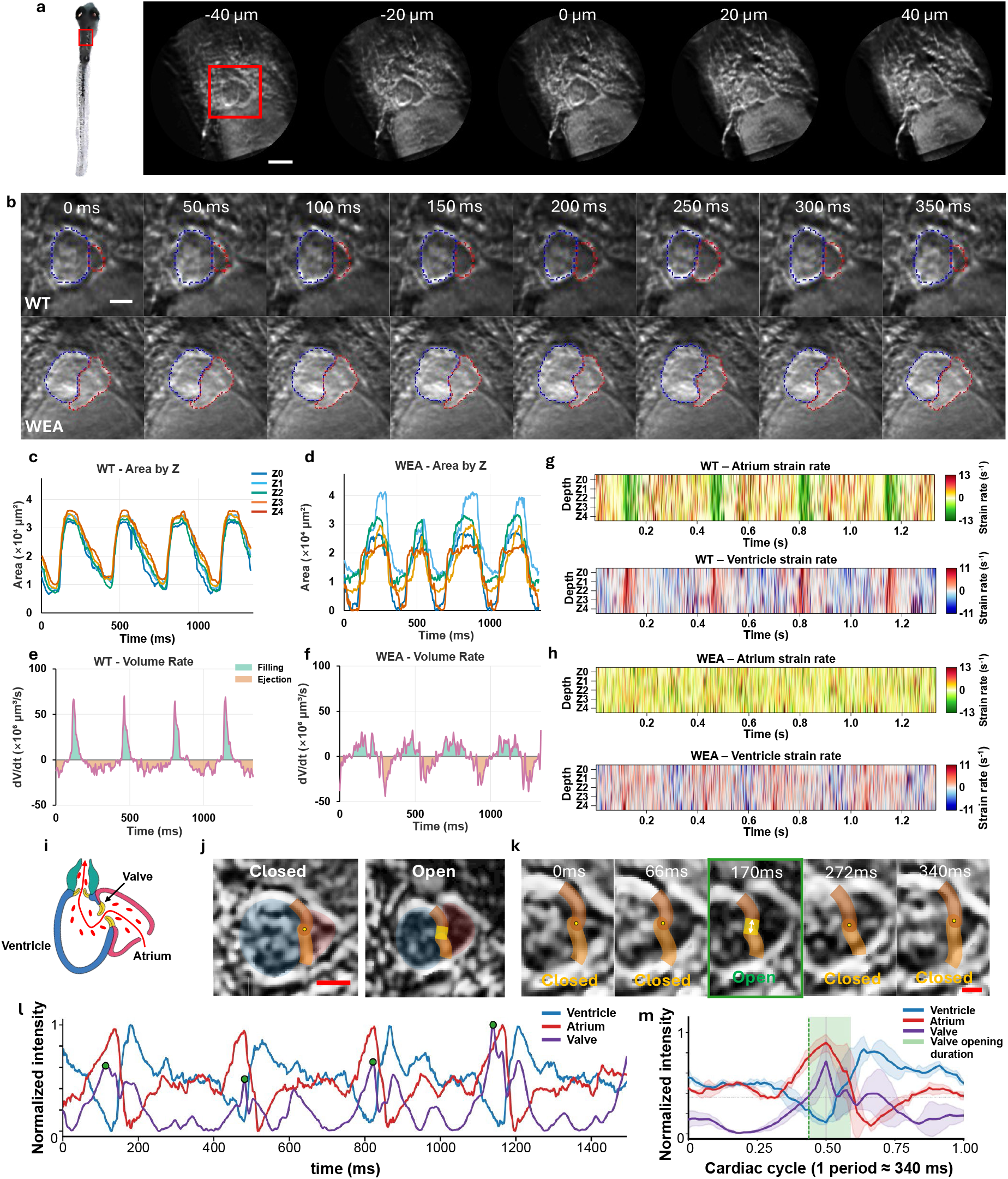
Label-free 4D imaging of cardiac and atrioventricular-valve dynamics in late-larval zebrafish. **a,** Label-free NIR-II SLIM images of the heart of a 20-dpf zebrafish acquired at 600 volumes *s*^−1^. Left, anatomical location of the imaged region; right, representative reconstructed sections spanning axial positions from −40 to 40 µm relative to the central cardiac plane (z = 0 here corresponds to z = −40 µm in Supplementary Video 1). Images in a and b are mirrored relative to Supplementary Videos 1 and 2. Scale bar, 200 µm. **b,** Representative cardiac dynamics over a 350-ms interval in wild-type (WT, top) and weak atrium (*wea*, *myh6*^−/−^; bottom) zebrafish. Dashed contours delineate the ventricle (blue) and atrium (red). Scale bar, 100 µm. **c, d,** Depth-resolved chamber-area dynamics across successive cardiac cycles in WT (**c**) and *wea* (**d**) hearts. Curves represent reconstructed axial planes at depths *Z*_0_ = −40 µm, *Z*_1_ = −20 µm, *Z*_2_ = 0 µm, *Z*_3_ = +20 µm, *Z*_4_ = +40 µm. **e, f,** Rate of ventricular volume change 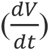 in WT (**e**) and *wea* (**f**) hearts, showing filling and ejection phases. **g, h,** Depth-resolved perimeter strain-rate maps for the atrium and ventricle in WT (**g**) and *wea* (**h**) hearts. WT hearts exhibit regular phase-locked deformation, whereas atrial deformation is markedly reduced in *wea* mutants. **i,** Schematic showing the anatomical location of the atrioventricular (AV) valve between the atrium and ventricle. **j,** Representative reconstructed images of the AV valve in closed and open states; colored overlays indicate the valve leaflets. Scale bar, 100 µm. **k,** Selected reconstructed volumes from a sequence acquired at 600 volumes s⁻¹ illustrate AV-valve opening and closure over one cardiac cycle. Scale bar, 50 µm. **l,** Simultaneously measured normalized ventricular, atrial, and valve signals over successive cardiac cycles; green markers indicate detected valve-opening events. **m,** Cycle-aligned ventricular, atrial, and valve dynamics over one cardiac period (∼340 ms). Green shading denotes the valve-opening interval.

Fish were imaged under label-free dark-field transillumination using a tungsten–halogen source spectrally filtered around 1,300 nm. Scattered light from the heart was collected by a custom imaging objective (Supplementary Fig. 1) and encoded into five subaperture views, enabling reconstruction of a complete cardiac volume from each camera exposure and thus continuous 4D imaging at 600 volumes s⁻¹ (Supplementary Fig. 6b).

NIR-II SLIM resolved the atrial and ventricular boundaries throughout the reconstructed axial range, allowing chamber morphology to be followed simultaneously across depth and time (Fig. 2a,b). Phase-resolved contours at the central depth captured the coordinated deformation of the two chambers over individual cardiac cycles (Fig. 2b). Depth-resolved cardiac dynamics in wild-type and *wea* zebrafish are shown in Supplementary Videos 1 and 2, respectively. We then segmented the atrium and ventricle throughout the volume using a U-Net model trained on manually annotated frames (Methods) and generated depth-resolved ventricular cross-sectional area–time traces (Fig. 2c,d). Integration of these segmented cross-sectional areas across depth yielded volumetric chamber measurements, from which ventricular volume-rate dynamics were quantified (Fig. 2e,f). Compared with wild-type hearts, *wea* hearts exhibited markedly altered ventricular filling dynamics and a reduced peak ventricular filling rate, consistent with the diminished atrial contribution expected from impaired atrial myosin function.

Beyond chamber-volume measurements, the volumetric data enabled depth-resolved characterization of myocardial deformation. Non-rigid registration was used to estimate contour displacement over time and derive perimeter strain-rate maps (fractional rate of change of the chamber perimeter, s^−1^; positive values indicate expansion) across the cardiac volume (Fig. 2g,h). Wild-type hearts exhibited regular, phase-locked deformation across depth, whereas *wea* hearts showed markedly reduced atrial deformation and less coordinated spatiotemporal dynamics. In contrast, ventricular perimeter strain rate was comparatively preserved, consistent with a predominantly atrial contractile defect in which ventricular function is affected only secondarily^40^.

We next examined whether the same imaging speed could resolve the substantially faster motion of the AV valve (Fig. 2i). Depth-resolved cardiac sections allowed us to identify the plane in which the valve leaflets were most clearly visualized and to distinguish their open and closed configurations (Fig. 2j). Consecutive reconstructed volumes captured valve opening and closure at 1.67-ms temporal intervals, directly resolving leaflet motion that would otherwise be blurred or undersampled at conventional volumetric imaging rates (Fig. 2k). We then aligned valve state with the atrial and ventricular dynamics over the cardiac cycle, revealing the temporal coordination between chamber contraction, filling, and valve opening (Fig. 2l). Cycle-aligned analysis further yielded the valve-opening profile and enabled extraction of its onset, duration, and termination (Fig. 2m). Across cycles, valve opening occupied approximately 15% (∼50 ms) of the ∼340-ms cardiac period, providing a quantitative readout of valve dynamics at a postembryonic stage that has not, to our knowledge, been accessible to high-speed *in vivo* imaging^42^.

Together, these results demonstrate that NIR-II SLIM extends high-speed cardiac microscopy into a postembryonic developmental window that is difficult to access with conventional visible-light imaging. By combining label-free NIR-II contrast with millisecond-scale volumetric acquisition, the method enables simultaneous quantification of chamber deformation, depth-dependent mechanics, and millisecond-scale AV-valve dynamics in intact, pigmented late-larval zebrafish.

### High-speed vascular and lymphatic imaging in mice *in vivo*

Blood and lymphatic vessels form 3D networks whose transport dynamics often extend beyond a conventional microscopic field of view (FOV). Volumetric imaging over an extended FOV is therefore important for maintaining connectivity across vessel branches, resolving vessels that overlap in 2D projections, and comparing upstream and downstream regions of interest (ROIs) within the same network. These capabilities are particularly relevant to tumor-associated lymphatics, in which tumor growth and lymph node involvement can alter vessel architecture, contractile transport, and drainage pathways^43,44^. NIR-II fluorescence is well suited to such measurements as reduced tissue scattering and autofluorescence at wavelengths above 1,000 nm improve visualization of subsurface vascular and lymphatic structures.

We used NIR-II SLIM to image blood flow in the mouse ear and lymphatic transport from a tumor in a xenograft-bearing mouse *in vivo*. The system was operated in epi-illumination with a 971-nm diode laser for excitation. A custom high-magnification objective was used for large-area mapping of the mouse ear vasculature by stitching adjacent volumetric tiles, whereas a custom low-magnification objective was used for dynamic imaging of selected ear ROIs and for lymphatic imaging (Supplementary Fig. 1). As the NIR-II fluorophore, we used Chrom7, a chromenylium heptamethine dye reported to be among the brightest members of its dye series^30^ (Fig. 3a and Methods). Micellar encapsulation enabled aqueous administration of Chrom7 for high-contrast visualization. In phosphate-buffered saline, Chrom7 micelles exhibited an absorption maximum at 992 nm and an emission maximum at 1,020 nm (Fig. 3b), with a long-wavelength emission tail detected between 1,100 and 1,300 nm.

**Fig. 3.**
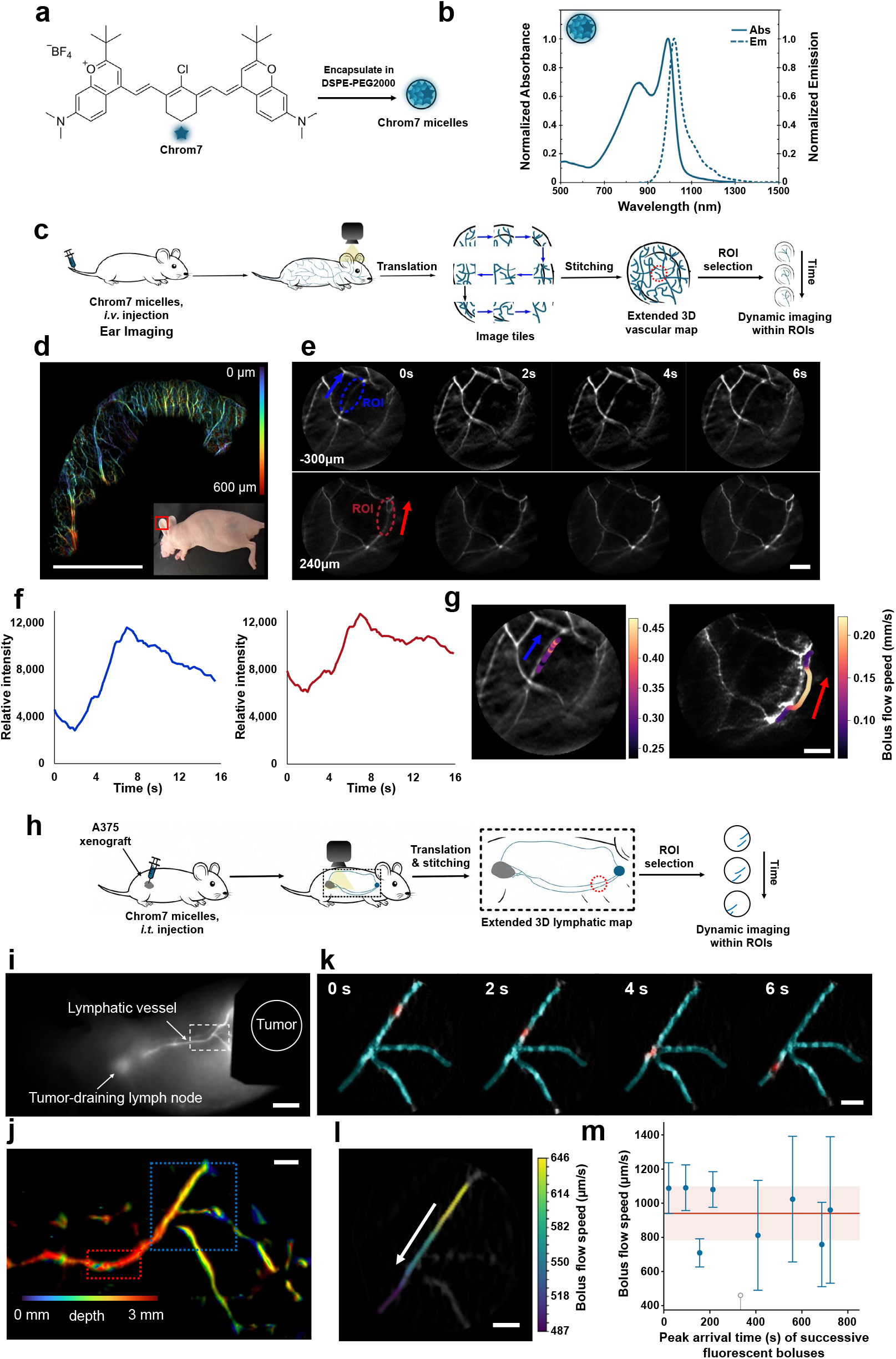
High-speed volumetric imaging of vascular and lymphatic transport *in vivo*. **a,** Chemical structure of Chrom7 and schematic of encapsulation in DSPE–PEG2000 micelles for aqueous administration. **b,** Normalized absorption (solid line) and emission (dashed line) spectra of Chrom7 micelles in phosphate-buffered saline. **c,** Workflow for extended-field vascular imaging. Following intravenous administration of Chrom7 micelles, adjacent volumetric fields are acquired by sample translation and stitched into an extended three-dimensional vascular map, from which regions of interest (ROIs) are selected for high-speed dynamic imaging; the resulting map and dynamic data are shown in d and e–g, respectively. **d,** Depth-coded map of mouse ear vasculature assembled from volumetric tiles acquired at 20 volumes s⁻¹. Color denotes reconstructed axial position over an approximately 600-µm depth range; inset shows the corresponding visible-light image and imaged region. Scale bar, 5 mm. **e,** Representative 20-volumes-s⁻¹ time series showing Chrom7 bolus passage through vascular ROIs located at different depths. Scale bar, 600 µm. **f,** Fluorescence-intensity traces from the ROIs in **e** (left, blue ROI at −300 µm; right, red ROI at +240 µm), resolving bolus arrival, peak intensity, and washout. **g,** Vessel centerlines in the reconstructed planes corresponding to the two ROIs in e, color-coded by spatially smoothed apparent bolus-propagation speed. Estimates in regions with insufficient local support were constrained by the corresponding ROI-level estimate. Scale bar, 600 µm. **h,** Workflow for imaging tumor-draining lymphatics in an A375 xenograft-bearing mouse. Following intratumoral injection of Chrom7 micelles, translation and stitching provide an extended three-dimensional map of the drainage pathway, followed by targeted high-speed imaging of selected ROIs (i–m). **i,** Macroscopic SWIR fluorescence image showing the tumor, tumor-draining lymph node, and connecting lymphatic vessel beneath the intact skin. Scale bar, 5 mm. **j,** Depth-coded map of the tumor-draining lymphatic network assembled from volumetric tiles acquired at 100 volumes s⁻¹ and denoised with DeepCAD-RT before reconstruction (Methods). Boxes indicate regions used for dynamic analysis. Scale bar, 1 mm. **k,** Representative time series acquired at 100 volumes s⁻¹, showing propagation of a discrete fluorescent bolus through the blue-box ROI in **j.** Camera-frame sequences used to generate the map and dynamic data in j–m were denoised with DeepCAD-RT before volumetric reconstruction (Methods). Scale bar, 600 µm**. l,** Three-dimensional trajectory of a representative fluorescent bolus, color-coded by its apparent propagation velocity. Scale bar, 600 µm. **m,** Pulse-to-pulse transport dynamics in the red-box ROI in **j**, quantified from a separate dataset acquired at 30 volumes s⁻¹. Each filled symbol shows the estimated apparent propagation speed of one fluorescent bolus, plotted at its peak arrival time; vertical bars indicate the 95% confidence interval of the estimate. The horizontal line and shaded band indicate the mean ± s.d. across the included boluses (n = 8, from one vessel in one mouse). The open symbol marks an additional bolus for which propagation speed could not be reliably estimated (R² < 0.5); this event was excluded from the speed summary. Variability in both arrival time and propagation velocity reveals irregular pulse-to-pulse lymphatic transport.

We first evaluated high-speed vascular imaging in the mouse ear, which provides a thin, optically accessible vascular bed containing arteries, veins, and capillary networks. Figure 3c illustrates the acquisition strategy used to combine large-area mapping with high-speed local measurements. Following intravenous injection of Chrom7 micelles, adjacent volumetric FOVs were acquired by free-form translation of the motorized stage, and subsequently registered and stitched to generate an extended 3D vascular map; the short per-volume acquisition minimized motion blur between positions. This approach preserves the depth information within each tile while expanding the lateral coverage beyond that of a single microscopic FOV. The resulting depth-coded reconstruction revealed vessels distributed across multiple axial planes, including subsurface branches that were poorly resolved in the corresponding visible-light image (Fig. 3d). The calibrated axial range is characterized in Supplementary Fig. 3, and the depth dependence of the *in vivo* signal-to-background ratio is shown in Supplementary Fig. 6c.

Guided by this large-area map, we then switched to the low-magnification objective and selected vascular ROIs for high-speed dynamic imaging at 20 volumes s⁻¹. Because each camera exposure encoded an entire volume, vessels at different depths were sampled simultaneously during the same bolus passage. We monitored Chrom7 fluorescence at ROIs distributed across different vessel segments and depths (Fig. 3e and Supplementary Videos 3 and 4). The corresponding intensity traces resolved bolus arrival, peak fluorescence, and the onset of washout at each location (Fig. 3f).

To quantify transport, we analyzed the first-pass rise in Chrom7 fluorescence along selected vessel centerlines in position–time maps. Figure 3g shows the resulting spatially smoothed apparent bolus-propagation speed profiles. The displayed estimates ranged from 0.23 to 0.43 mm s⁻¹ in the blue ROI and from 0.09 to 0.21 mm s⁻¹ in the red ROI. For context, intravital line-scan measurements in mouse ear dermis reported red-blood-cell (RBC) velocities above 0.1 mm s⁻¹ in arterioles and below 0.1 mm s⁻¹ in capillaries and venules under ketamine–xylazine anesthesia^45^. Third-harmonic-generation (THG) line scanning under fentanyl–medetomidine–midazolam anesthesia also revealed strongly time-dependent arteriolar velocities, with transient systolic maxima ranging from 1.7 to 9.4 mm s⁻¹ in representative recordings^46^. These studies provide physiological context rather than direct validation, because the present measurement tracks a fluorescent-bolus wavefront rather than individual RBCs, and vessel characteristics, temporal sampling, and anesthetic conditions differ. We therefore interpret the measured values as apparent bolus-propagation velocities and local proxies for blood-flow speed. Depth-resolved reconstruction enabled the selected vascular regions to be analyzed separately at their respective axial positions. Using the high-magnification objective, NIR-II SLIM also resolved the three-dimensional vascular architecture of the mouse hind paw through intact skin (Supplementary Fig. 7 and Supplementary Video 5).

We next applied the same strategy to lymphatic transport in an A375 tumor-bearing mouse (Fig. 3h). Following local intratumoral injection of Chrom7 micelles, a macroscopic short-wave infrared (SWIR) image was first acquired to identify the fluorescent drainage pathway beneath the intact skin (Fig. 3i). We then translated the sample across this pathway and stitched the reconstructed subvolumes to obtain an extended 3D map of the tumor-draining lymphatic network (Fig. 3j). The macroscopic SWIR image established the overall drainage route, whereas NIR-II SLIM provided depth-resolved visualization and enabled selected regions to be interrogated dynamically at high speed.

High-speed imaging revealed that lymphatic transport occurred as discrete fluorescent boluses rather than as a steady signal (Supplementary Videos 6 and 7). To quantify these events, we analyzed the 100-volumes-s⁻¹ recording from the blue-box ROI in Fig. 3j. For each pair of consecutive volumes, the preceding volume was subtracted from the current volume, and the advancing positive-intensity peak was tracked along the segmented 3D vessel centerline. We treated this peak as the leading edge of the fluorescent bolus. Its position as a function of time defined the wavefront trajectory (Fig. 3k), and its frame-to-frame displacement divided by the intervolume interval yielded the apparent lymph-bolus propagation velocity (Fig. 3l). This quantity represents the propagation velocity of the fluorescent boluses and serves as a proxy for quantifying lymphatic transport. Depth-dependent SBR and the effect of denoising are characterized in Supplementary Fig. 6a, d.

Using a separate dataset acquired at 30 volumes s⁻¹, we analyzed successive pulse-like events in the red-box region of Fig. 3j to characterize pulse-to-pulse transport dynamics (Fig. 3m). The horizontal line and shaded band indicate the mean ± s.d. across boluses (n = 8 boluses from one tumor-draining vessel in one mouse); the open symbol marks one additional bolus whose propagation speed could not be reliably estimated (R² < 0.5) and was excluded from the speed summary. Both the arrival times and propagation velocities varied markedly among successive fluorescent boluses, revealing temporal heterogeneity that would be obscured by static or time-averaged lymphangiography. The inter-pulse intervals were highly irregular, with a coefficient of variation of approximately 0.45, substantially greater than the contraction-interval variability reported for normal rat mesenteric collecting lymphatics (∼0.02)^47,48^, although species, vessel bed, and measured quantity differ. This elevated variability may reflect less regular lymphatic transport within the tumor-draining vessel. Together, these measurements demonstrate that high-speed volumetric imaging can resolve not only the trajectories of lymphatic transport, but also its pulsatility and pulse-to-pulse variability.

Overall, these vascular and lymphatic experiments demonstrate how NIR-II SLIM combines complementary spatial and temporal scales: tiled volumetric imaging provides 3D network context, while targeted high-speed acquisition resolves rapid local transport dynamics. This combination enables propagation speed, pulsatility, and transport variability to be quantified within the anatomical context of the surrounding 3D vascular or lymphatic network.

## Discussion

### Multiplexing advantage under detector-noise-limited conditions

InGaAs image sensors generally exhibit substantially higher read noise and dark-current noise than silicon-based cameras^25^, making detector noise a major limitation in NIR-II fluorescence imaging, particularly at low photon levels. NIR-II SLIM partially mitigates this limitation through optical compression. By squeezing each perspective image along one spatial dimension, the system concentrates the optical signal onto fewer detector pixels and therefore reduces the number of noisy pixel readouts contributing to each reconstructed volume.

Under detector-noise-limited conditions, combining information optically before detection can yield a higher SNR than measuring the same information independently across a larger number of detector elements^28^. In our system, the InGaAs sensor operates in a read-noise-dominated regime at these frame rates, so concentrating the signal onto fewer effective pixels reduces the number of read-noise contributions per reconstructed voxel (Supplementary Note 2); this advantage does not hold in the shot-noise-limited regime. For an ideal compression factor *N*, with conserved photon throughput, independent and identically distributed pixel noise, and a well-conditioned reconstruction, the SNR improvement can approach √*N*. Here, *N* corresponds approximately to the anamorphic compression ratio along the squeezed axis relative to the unsqueezed image dimension. Numerical simulations illustrating this SNR improvement are provided in Supplementary Note 2, Supplementary Fig. 10 and Supplementary Table 1. Thus, optical squeezing not only accommodates multiple perspective views on a limited-format InGaAs sensor but also improves detection sensitivity under conditions commonly encountered in NIR-II imaging.

### Diverse sampling of 3D spatial-frequency space

Light-field reconstruction can be interpreted as a limited-angle tomographic inverse problem. Rather than sequentially sampling the object at different spatial positions, as in confocal microscopy, or acquiring a series of physical planes, as in light-sheet microscopy, light-field imaging records multiple angular projections simultaneously. Under the Fourier-slice interpretation, each perspective view contributes a 2D region of support within the 3D spatial-frequency domain. The orientations and extents of these Fourier slices determine the coverage of the object’s spatial-frequency spectrum and thereby influence the resolution, isotropy, and conditioning of the 3D reconstruction.

Anamorphic compression modifies this frequency-space support. Compressing a perspective image along one spatial axis reduces its recoverable bandwidth along the corresponding image-space direction, transforming the approximately circular frequency support of an unsqueezed view into an anisotropic, elliptical support. If all perspective images were compressed along the same object-space direction, the measurements would provide substantially weaker disparity information and spatial-frequency coverage along that axis. This anisotropy would worsen the conditioning of the inverse problem and lead to direction-dependent loss of high-frequency features (Supplementary Note 1).

The Dove-prism rotations in NIR-II SLIM are therefore essential for preserving complementary spatial-frequency information. By rotating the unsqueezed, high-bandwidth axis of each view to a different orientation, the system distributes the anisotropic frequency supports across the lateral frequency plane. Their combined coverage recovers substantially broader and more isotropic lateral spatial bandwidth than would be available from identically oriented compressed views. This sampling diversity improves the reconstruction of fine structures, reduces orientation-dependent artifacts, and allows SLIM to retain much of the lateral spatial information of unsqueezed light-field measurements despite strong anamorphic compression (Supplementary Note 1).

### Limited-angle reconstruction and opportunities for resolution enhancement

Despite its improved frequency-space coverage, NIR-II SLIM remains subject to the fundamental limitations of sparse-view and limited-angle tomography. The finite number of subaperture views and restricted angular range leave portions of the 3D spatial-frequency spectrum weakly sampled or unmeasured. This missing-cone problem primarily limits axial-frequency support, producing lower axial than lateral resolution and increasing sensitivity to noise and model mismatch. Physics-informed reconstruction can suppress some associated artifacts by enforcing consistency with the calibrated optical model, but it cannot fully recover spatial frequencies that are absent from the measurements without introducing additional information or prior assumptions.

One potential strategy for expanding the accessible frequency support is to combine NIR-II SLIM with structured illumination. Multiplying the object by a spatially modulated illumination pattern shifts otherwise inaccessible object frequencies into the detection passband. By acquiring patterns with multiple phases and orientations, the shifted spectra could supplement the anisotropic supports provided by the subaperture views, extend both lateral and axial frequency coverage, and improve the reconstruction of high-frequency features. Appropriately designed illumination patterns may also reduce resolution anisotropy and partially fill the missing cone.

These gains would come with important tradeoffs. Structured illumination requires multiple exposures per reconstructed volume, reducing the effective volumetric imaging rate and increasing sensitivity to motion between patterns. Additional exposures may also increase illumination dose and complicate synchronization, calibration, and reconstruction. Thus, structured-illumination NIR-II SLIM would be most valuable for applications in which enhanced and more isotropic spatial resolution is more important than the maximum snapshot acquisition speed. Adaptive pattern selection, high-speed spatial light modulators, and joint physics-informed reconstruction may help reduce the number of required measurements and balance spatial-frequency coverage against temporal resolution.

### Out-of-focus light and optical sectioning

Similar to conventional light-field microscopy, NIR-II SLIM does not intrinsically reject out-of-focus light^49^. Although computational reconstruction can refocus the measurements and deconvolve the 3D volume, photons originating outside the reconstructed plane are still detected and contribute background and shot noise. This limitation becomes increasingly important in thick or densely labeled specimens, where accumulated out-of-focus fluorescence can reduce contrast and obscure weak structures.

One potential remedy is targeted illumination^50,51^, in which excitation is restricted to selected structures or regions of interest identified by a preliminary wide-field or fluorescence measurement. By illuminating only the relevant volume, targeted illumination suppresses background generation at its source and improves photon efficiency. Another possibility is random-illumination microscopy^52,53^, which uses the statistical fluctuations of laser-speckle illumination to distinguish in-focus signals from spatially uncorrelated out-of-focus background. Combining these illumination strategies with SLIM could provide computational optical sectioning while retaining substantially higher volumetric imaging speeds than conventional point-or plane-scanning methods.

### Quantification of flow speed

NIR-II SLIM can resolve the 3D propagation of contrast-agent signals through vascular or lymphatic networks. However, estimating flow speed from the leading edge of an injected bolus may be unreliable because the measured bolus dynamics are influenced by injection conditions, dispersion, vessel geometry, tracer diffusion, and temporal variations in concentration. The apparent propagation velocity may therefore differ from the local fluid velocity, particularly in branching networks or vessels with pulsatile or recirculating flow.

A promising solution is to combine NIR-II SLIM with laser-speckle contrast imaging^54^. Fluorescence imaging could first localize vessels and recover their 3D geometry, while coherent illumination could then be directed to selected vessels to measure speckle fluctuations associated with blood motion. This multimodal approach would combine depth-resolved anatomical information with flow-sensitive contrast, enabling more reliable estimation of relative or absolute flow velocities. Accurate quantification would require calibration against vessel diameter, exposure time, scattering conditions, and an appropriate speckle-flow model.

### Optical-diffusion limit

Despite the reduced scattering achieved at longer wavelengths, NIR-II SLIM remains primarily dependent on ballistic and weakly scattered photons that preserve sufficient directional information for light-field reconstruction. As imaging depth increases, multiple scattering progressively randomizes photon directions and destroys the angular disparities required for accurate tomographic reconstruction. Consequently, the penetration depth of NIR-II SLIM remains fundamentally limited by the onset of optical diffusion.

*In vivo* optical clearing may provide a route to extending this depth range. Recent studies have shown that strongly absorbing dyes, including the food additive tartrazine, can transiently reduce tissue scattering by modifying the tissue refractive index environment^55^. Applying such agents before imaging could increase photon transmission, preserve angular information over greater depths, and improve the signal-to-background ratio of NIR-II SLIM. Future work should determine whether this approach is compatible with NIR-II wavelengths and evaluate its clearing efficiency, onset and recovery times, systemic effects, and influence on physiological measurements.

## Methods

### Optical design and system setup

For NIR-II SLIM operated in dark-field trans-illumination mode, we used a custom-designed illumination lens (Supplementary Fig. 1) and objective lens (Supplementary Fig. 1; 5.2× magnification with a 200-mm tube lens; numerical aperture, NA = 0.44). Illumination was provided by a halogen lamp spectrally filtered with a bandpass filter centered at approximately 1,300 nm (bandwidth, 25 nm).

On the detection side, the back aperture of the objective was partitioned into a linear array of five subapertures, each forming an independent perspective view through a Dove prism and two lenslets. In each channel, a Dove prism (PS990, Thorlabs) was positioned immediately before the first lenslet (aperture, 5 mm; focal length, 127 mm; Sunday Optics). The five Dove prisms were oriented such that the corresponding perspective images were rotated by angles evenly spanning 0° to 180°, with the image rotation in each channel equal to twice the rotation angle of the Dove prism. An identical second lenslet was placed at the intermediate image plane and served as a field lens. This field lens redirected the off-axis chief rays and rendered the individual channels approximately telecentric in image space, thereby minimizing vignetting across the five perspective views. Each subaperture channel produced a rotated perspective image with a magnification of approximately 3.0× and an effective NA of 0.065. The five Dove-prism/lenslet channels were mounted in custom 3D-printed mechanical holders (Supplementary Fig. 2).

The five perspective images were subsequently relayed to the detector through an anamorphic optical system consisting of a spherical achromatic doublet (88-598, Edmund Optics) and two orthogonally oriented cylindrical-lens assemblies. The x-oriented assembly incorporated a 500-mm-focal-length cylindrical lens (LJ1144L1-C, Thorlabs), whereas the y-oriented assembly comprised two 150-mm-focal-length cylindrical lenses (LJ1629L1-C, Thorlabs). The back focal planes of the two cylindrical-lens assemblies were co-located, producing anisotropic magnification along the two orthogonal spatial axes (Supplementary Fig. 2). This anisotropic relay compressed the perspective-image array along one dimension while preserving spatial sampling along the orthogonal dimension. The resulting image array was recorded using a cooled InGaAs camera (C-RED 2, First Light Imaging; 640 × 512 pixels; sensor temperature, −40 °C; readout noise <30 e⁻ r.m.s. in correlated-double-sampling mode at 600 frames s⁻¹ and dark current ≈300 e⁻ pixel⁻¹ s⁻¹ per the manufacturer’s characterization^56^). At the 600 volumes s⁻¹ used here (≈1.67-ms exposure), dark charge per frame is <1 e⁻, so readout noise dominates the detector noise. The Dove-prism rotations generated differently oriented perspective images, providing complementary angular sampling in Fourier space.

For NIR-II SLIM operated in widefield epi-fluorescence mode, two custom objective lenses, designed in-house and assembled from catalogue achromatic elements (prescriptions and part numbers in Supplementary Fig. 1), were used to provide different combinations of spatial resolution and FOV. The higher-magnification objective was the same 5.2×, NA 0.44 objective used for dark-field imaging (working distance 6.18 mm; elements AR-coated for 1,050–1,700 nm) and was used for large-area mapping of the mouse ear vasculature and for mouse paw imaging. A lower-magnification objective (Supplementary Fig. 1; 1.9× magnification with a 200-mm tube lens; NA = 0.16; working distance 109.89 mm) provided a larger FOV and was used for dynamic imaging of mouse ear ROIs and for lymphatic imaging.

Fluorescence excitation was provided by a laser diode centered at approximately 971 nm (model L960H1, Thorlabs), together with neutral-density and spatial-filtering optics. The fluorescence detection path used the same subaperture encoding and anamorphic relay optics described above. An additional emission filter set (1,100-nm long-pass filter and 1,300-nm short-pass filter) was placed in the detection path to suppress residual excitation light before the fluorescence signal reached the InGaAs camera.

### System calibration, forward model, and reconstruction

For system calibration, a pinhole array (R1L3S5P, Thorlabs) was axially scanned through the imaging volume. For the higher-magnification objective, we used a pinhole pattern with a 100-µm pitch and 20-µm-diameter apertures and acquired calibration images at 30-µm axial intervals over a total range of 600 µm (±300 µm). For the lower-magnification objective, a pattern with a 400-µm pitch and 80-µm-diameter apertures was used, with calibration images acquired at 150-µm axial intervals over a total range of 3 mm (±1.5 mm). PSFs at intermediate depths were obtained by interpolation of the calibrated PSFs. For the main-figure datasets, axial reconstruction spacing was 20 µm for zebrafish cardiac imaging and mouse ear vasculature mapping, and 60 µm for mouse ear dynamic vascular imaging and tumor-draining lymphatic imaging. Mouse paw data were reconstructed at 20-µm axial spacing. The depth-dependence datasets in Supplementary Fig. 6 were reconstructed at 12-µm (high-magnification) and 60-µm (low-magnification) axial spacing. High-magnification volumes covered a 600-µm axial range, whereas low-magnification (mouse ear dynamic and lymphatic) volumes covered a 3-mm axial range. At each axial position, the PSFs of all five subaperture views were recorded.

The depth-resolved PSF stack for each subaperture view was used to construct a view-specific projection operator, ***P_i_***. The depth-dependent lateral disparity of each view was determined by linear fitting of the measured PSF positions as a function of axial depth and was used to interpolate the PSFs at intermediate axial positions. In parallel, a view-specific affine transformation, ***T_i_***, was estimated relative to the central subaperture to account for fixed image rotation, anisotropic magnification, and other camera-plane geometric distortions. For calibration and visualization, the measured PSFs were rectified into a common intermediate-image coordinate system using 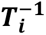, while preserving the depth-dependent disparities required for 3D reconstruction.

The forward model for the *i*-th subaperture view was therefore expressed as *I_i_* = **T*_i_*P*_i_****ρ* + *η_i_*, where *I_i_* denotes the compressed camera measurement from the *i*-th perspective view, *ρ* represents the 3D object distribution, and *η_i_* represents measurement noise. During calibration, *ρ* corresponds to the known 3D pinhole distribution. The complete multi-view forward operator was constructed by combining the five view-specific operators and was subsequently used for iterative Richardson–Lucy deconvolution and 3D reconstruction. Reconstruction was performed in camera coordinates using the combined operators **T*_i_*P*_i_*** rather than on rectified views, with the standard multiplicative Richardson–Lucy update, a uniform initial volume, and a fixed stopping criterion of 8 iterations.

For typical datasets, 5–10 iterations were sufficient to obtain stable reconstructions. The reconstruction algorithm was implemented in MATLAB with GPU acceleration (NVIDIA GeForce RTX 4090) and required approximately 0.5 s per volume (∼2 volumes s^−1^) for offline reconstruction, excluding data transfer. Acquisition at up to 600 volumes s^−1^ is thus decoupled from reconstruction, which is performed post hoc.

### NIR-II fluorophore micelle preparation

Chrom7 was synthesized as previously described^30^. For *in vivo* imaging, Chrom7 was encapsulated within DSPE-PEG2000 micelles as follows: A 6 mg/mL solution of 18:0 PEG2000PE (Avanti Polar Lipids) was prepared in MilliQ water (2 mL). Chrom7 (0.1 mg) was dissolved in dimethyl sulfoxide (DMSO) (1 mL) and added to the surfactant solution with thorough mixing. The mixture was then transferred to a 15 mL conical tube and probe-sonicated on ice (3 min, 35% amplitude) using a QSonica (Q125) probe sonicator. The resulting dispersion was concentrated and washed using a 10 kDa molecular weight cutoff centrifugal filter (Amicon Ultra-15) at 4,000 g for 20 min. Buffer exchange into phosphate-buffered saline (PBS; 3 mL) was performed by repeated centrifugation (three cycles total) to reduce the DMSO content to <1%. After filtration through a 0.22-µm syringe filter, micelles were analyzed by dynamic light scattering using a Malvern Zetasizer Nano instrument. Absorption and emission of the micelles were measured on the JASCO V-770 UV-Visible/NIR spectrophotometer and the Horiba Fluorometer PTI QM-400 with a liquid nitrogen cooled InGaAs detector (Horiba Edison DSS IGA 020L), respectively.

### Animal preparation and imaging

All animal experiments were conducted in accordance with the University of California, Los Angeles guidelines and approved by the Animal Research Committee (Protocol numbers: ARC-2021-130, ARC-2018-047, and ARC-2024-086). Female homozygous athymic NU/J mice (6–16 weeks old, 20–25 g; The Jackson Laboratory) were used for whole-body imaging studies. Mice were anesthetized with 1–4% isoflurane. For intravenous (*i.v.*) administration, a catheter was prepared using a 29-gauge needle (VetriJec) connected to polyethylene tubing, flushed with isotonic saline, inserted into the tail vein, and secured with tissue adhesive. Micelle-encapsulated Chrom7 was administered via the catheter using a 29-gauge syringe followed by a saline flush. All *i.v.* doses totaled 200 µL (62 nmol) per mouse. For intratumoral (*i.t.*) administration, micelle-encapsulated Chrom7 was directly administered using a 29-gauge syringe. All *i.t.* doses were 20–40 µL (6.2–12.4 nmol) per tumor. All solutions were filtered through a 0.22-µm syringe filter before use. Mice were imaged post-administration and subsequently euthanized by isoflurane overdose followed by cervical dislocation.

Xenograft procedures: A375 cells were obtained from the American Type Culture Collection and maintained in Dulbecco’s Modified Eagle Medium (DMEM; Life Technologies, 11995073) supplemented with 10% fetal bovine serum, 1 mM sodium pyruvate, and 1% penicillin–streptomycin at 37 °C in 5% CO₂. Cells were cultured to 70–80% confluency, harvested, counted, and resuspended in sterile PBS at a concentration of 1.5–2 × 10^7^ cells per mL. For xenograft implantation, the resuspended cells (100 µL, 1.5– 2 × 10^6^) were mixed 1:1 with Matrigel (100 µL). Mice were anesthetized with 2–4% isoflurane, and the cell suspension was injected subcutaneously into the hind flank using a 25-gauge syringe. The injection site was held for ∼30 s before needle withdrawal. Tumor growth was monitored daily by caliper measurements.

### Zebrafish cardiac imaging and heart chamber analysis

Wild-type (AB strain) and *wea* (*myh6*^−/−^) zebrafish at 15–25 days post-fertilization (dpf) were briefly anesthetized with 0.016% MS-222 (tricaine methanesulfonate) and embedded in 0.5% low-melting-point agarose in a glass-bottom dish. The heart was positioned within the working distance of the objective, and imaging was performed using dark-field transillumination centered at 1,300 nm. Heart rate and anesthesia depth were monitored throughout the imaging session.

Four-dimensional cardiac datasets were acquired at 600 volumes s⁻¹. Bulk specimen motion was corrected by 3D image registration using SimpleElastix. The atrium and ventricle were manually annotated in representative frames and segmented using separate U-Net models. Image intensity and normalized axial position were provided as inputs, and the networks were trained using a combined Dice–cross-entropy loss. Predicted masks were subjected to morphological cleanup and restricted to the largest connected component. Atrial and ventricular volumes were calculated independently by integrating the segmented cross-sectional areas along the axial dimension. For Fig. 2g,h, chamber deformation at each analyzed depth was quantified using global perimeter strain rate, defined as the temporal rate of change of the logarithm of the segmented chamber perimeter. Positive values indicate expansion.

### Zebrafish heart valve analysis

High-speed valve sequences were normalized to the 0.5–99.5th intensity percentiles and background-subtracted using the per-pixel temporal median; the valve-openness signal was smoothed with a Hilbert-envelope filter prior to event detection. An operator-defined line crossing the atrioventricular valve was used to generate an M-mode kymograph. Leaflet position was identified from the local intensity extremum with subpixel interpolation, and projected displacement and velocity were calculated using the calibrated pixel size and frame rate. Leaflet masks were manually annotated in selected key frames and propagated only across short intervals for which the interpolated boundaries agreed with the images.

Valve opening was quantified from leaflet separation or the projected area of the orifice between the leaflets. A fixed-ROI darkening signal, calculated relative to the per-pixel temporal median, was used only as an auxiliary open-state proxy. Candidate opening events were searched within cardiac cycles defined from the atrial and ventricular signals without enforcing one event per cycle. Phase averaging was used only to display the mean valve cycle and was not used for beat-to-beat statistical measurements.

### Mouse blood and lymph vessel imaging and flow speed analysis

Mice were anesthetized with isoflurane in an induction chamber and subsequently positioned on a custom imaging holder equipped with a heated stage (model 69027, Thermostar). Anesthesia was maintained through a nose cone connected to the isoflurane delivery system, and respiration and overall physiological condition were monitored throughout the imaging sessions. For mouse ear vascular imaging, the ear was gently flattened and secured to a custom 3D-printed holder using surgical adhesive. Chrom7 micelles were administered intravenously through the tail vein. During imaging, the specimen was translated using a motorized XYZ stage (MCM3001m, Thorlabs) to acquire adjacent FOVs for large-area volumetric mapping. For the experiments shown in Fig. 3, mouse ear mapping (high-magnification objective) and dynamic imaging (low-magnification objective) were performed at 20 volumes s⁻¹. Lymphatic mapping and representative bolus dynamics in Fig. 3j–l were acquired at 100 volumes s⁻¹, whereas the separate dataset used for pulse-to-pulse analysis in Fig. 3m was acquired at 30 volumes s⁻¹.

For mouse ear vascular analysis, vessel centerlines and adjacent background regions were manually defined. Frames affected by bulk or respiratory motion were identified using a robust median-absolute-deviation threshold and excluded from quantitative analysis. Bolus-propagation speed profiles were derived from the temporal progression of fluorescence along vessel centerlines traced in the reconstructed planes. Where local estimates were insufficiently supported, they were regularized toward or replaced by the corresponding ROI-level estimate. The profiles were spatially smoothed for display, and the ranges reported for Fig. 3g refer to these displayed estimates. Quantitative fits used directly observed, non-excluded frames.

For lymphatic-vessel analysis, consecutive reconstructed volumes were temporally differenced, and the advancing positive-intensity peak was tracked along the manually traced three-dimensional vessel centerline. Apparent frame-to-frame bolus-propagation velocity was calculated from changes in the tracked centerline position divided by the intervolume interval. These measurements were interpreted as the propagation dynamics of fluorescent boluses and not as direct measurements of cross-sectional mean lymph-flow velocity or lymphatic contraction-wave speed.

Mouse paw vascular imaging: Male CD1 mice (2 months old) were provided by C.Y.K. under protocol ARC-2024-086. Anesthesia, physiological monitoring, and intravenous administration of Chrom7 micelles were performed as described above for mouse ear imaging. The hind paw was gently secured to the custom holder with surgical adhesive, and volumetric imaging was performed with the high-magnification objective (Supplementary Fig. 1) at 30 volumes s⁻¹ (Supplementary Fig. 7 and Supplementary Video 5).

Self-supervised denoising. Camera-frame sequences used for the lymphatic dynamic analyses in Fig. 3k,l (100 volumes s⁻¹) and Fig. 3m (30 volumes s⁻¹) were denoised with DeepCAD-RT^57,58^ before volumetric reconstruction, as were the recordings used for the denoising comparison in Supplementary Fig. 6d. Mouse ear mapping and dynamic recordings acquired at 20 volumes s⁻¹, label-free dark-field zebrafish recordings, and mouse paw recordings were reconstructed without DeepCAD-RT denoising. The same denoising pipeline was applied to the camera-frame sequences used to generate the lymphatic map in Fig. 3j. Local modifications to the code were limited to output-path handling, leaving the network architecture, loss function, and training procedure unchanged. Consistent with its self-supervised design, a separate model was trained on each recording (5 epochs, learning rate 2 × 10⁻⁵, batch size 1, Adam with β₁ = 0.5 and β₂ = 0.999, 16 feature maps, 150 × 150 × 150-voxel patches in x, y, and t with gaps of 90, 90, and 22–40 voxels, overlap factor 0.4); inference used the epoch-5 checkpoint with the same patch settings. Signal-to-background ratios in Supplementary Fig. 6a–c were computed from non-denoised reconstructions. Denoising was run in Python 3.14.3 with PyTorch 2.11.0 (CUDA 12.8) on an NVIDIA RTX 4090.

### Mouse lymph statistical analysis

Pulse events were mapped to their original acquisition times before statistical analysis. Pulse-event frequency and inter-event intervals were calculated separately within each continuously observed acquisition window, and intervals spanning gaps in observation were excluded. For each detected event, we quantified the bolus-propagation speed, transport distance, and longitudinal full width at half maximum of the fluorescent bolus.

The dataset used for Fig. 3m was acquired at 30 volumes s⁻¹, and event times and propagation speeds were calculated using this acquisition rate. Summary statistics describe variability among eight included bolus events within one vessel from one mouse and do not represent between-animal variability.

### Use of large language models

Claude (Anthropic) and ChatGPT (OpenAI) were used to assist with language editing of the manuscript and with debugging analysis and plotting scripts. No figures or images were generated with these tools. All outputs were reviewed, tested, and edited by the authors, who take full responsibility for the content.

## Data availability

Selected raw imaging datasets, calibration files, representative reconstructed volumes, and numerical source data are available at Zenodo (DOI: 10.5281/zenodo.22184824). The repository README specifies the files associated with each figure. Owing to their large size, the full raw tile sequences used to generate the stitched mouse ear map in Fig. 3d are not included in the public deposit and are available from the corresponding author upon reasonable request. Source data are provided with this paper.

## Code availability

The reconstruction algorithms and analysis codes are publicly available on GitHub at https://github.com/plapenda1996/NIR_II_SLIM_Reconstruction

## Contributions

D.Y.K., Z.Z., and L.G. conceived the idea. D.Y.K. and Z.Z. constructed the microscopes and developed the reconstruction algorithm, with assistance from R.Z. D.Y.K., Z.Z., E.M.S., and E.Y.L. conceived the mouse experiments. E.Y.L. prepared the Chrom7 micelles, established the xenografts, and performed the mouse surgical and dye-administration procedures. D.Y.K. and Z.Z. bred the zebrafish, and J.W. prepared the weak atrium (*wea*) zebrafish. C.Y.K. provided the CD1 mice and contributed to the paw vascular imaging experiments. D.Y.K. and Z.Z. acquired, processed, and analyzed the imaging data. T.K.H. supervised the zebrafish studies. E.M.S. supervised the mouse studies, along with the Chrom7 synthesis and micelle preparation. L.G. supervised the project. D.Y.K., Z.Z., and L.G. wrote the paper with input from all authors, and all authors reviewed and edited the manuscript.

## Acknowledgements

This work was supported by the National Institutes of Health (R01HL165318, RF1NS128488, and R35GM128761) and the U.S. Department of Energy (DE-SC0025928). We thank E. Mobley (Sletten laboratory) for assistance with xenograft implantation.

## References

1. Wang, T., Chen, Y., Wang, B., Gao, X. & Wu, M. Recent progress in second near-infrared (NIR-II) fluorescence imaging in cancer. Biomolecules 12, 1044 (2022).

2. Li, C., Chen, G., Zhang, Y., Wu, F. & Wang, Q. Advanced fluorescence imaging technology in the near-infrared-II window for biomedical applications. J. Am. Chem. Soc. 142, 14789–14804 (2020).

3. Miao, Y. et al. Recent progress in fluorescence imaging of the near-infrared II window. ChemBioChem 19, 2522–2541 (2018).

4. Roblyer, D. et al. Review of shortwave infrared imaging and spectroscopy in tissue [Invited]. Biomed. Opt. Express 16, 5028–5062 (2025).

5. Ding, F., Zhan, Y., Lu, X. & Sun, Y. Recent advances in near-infrared II fluorophores for multifunctional biomedical imaging. Chem. Sci. 9, 4370–4380 (2018).

6. Chen, Y., Xue, L., Zhu, Q., Feng, Y. & Wu, M. Recent advances in second near-infrared region (NIR-II) fluorophores and biomedical applications. Front. Chem. 9, 750404 (2021).

7. Tang, Y., Pei, F., Lu, X., Fan, Q. & Huang, W. Recent advances on activatable NIR-II fluorescence probes for biomedical imaging. Adv. Opt. Mater. 7, 1900917 (2019).

8. Jain, A., Homayoun, A., Bannister, C. W. & Yum, K. Single-walled carbon nanotubes as near-infrared optical biosensors for life sciences and biomedicine. Biotechnol. J. 10, 447–459 (2015).

9. Su, Y., Yu, B., Wang, S., Cong, H. & Shen, Y. NIR-II bioimaging of small organic molecule. Biomaterials 271, 120717 (2021).

10. Yu, Z., Eich, C. & Cruz, L. J. Recent advances in rare-earth-doped nanoparticles for NIR-II imaging and cancer theranostics. Front. Chem. 8, 496 (2020).

11. Chen, L. L., Zhao, L., Wang, Z. G., Liu, S. L. & Pang, D. W. Near-infrared-II quantum dots for in vivo imaging and cancer therapy. Small 18, 2104567 (2022).

12. Mu, J., et al. The chemistry of organic contrast agents in the NIR-II window. Angew. Chem. Int. Ed. 61, e202114722 (2022).

13. Huang, L.-Y., Zhu, S., Cui, R. & Zhang, M. Noninvasive in vivo imaging in the second near-infrared window by inorganic nanoparticle-based fluorescent probes. Anal. Chem. 92, 535–542 (2020).

14. Taber, L. A. Mechanical aspects of cardiac development. Prog. Biophys. Mol. Biol. 69, 237–255 (1998).

15. Lindsey, S. E., Butcher, J. T. & Yalcin, H. C. Mechanical regulation of cardiac development. Front. Physiol. 5, 318 (2014).

16. Cai, Z. et al. NIR-II fluorescence microscopic imaging of cortical vasculature in non-human primates. Theranostics 10, 4265–4276 (2020).

17. Zubkovs, V. et al. Spinning-disc confocal microscopy in the second near-infrared window (NIR-II). Sci. Rep. 8, 13770 (2018).

18. Wang, F. et al. Light-sheet microscopy in the near-infrared II window. Nat. Methods 16, 545–552 (2019).

19. Wang, F., et al. In vivo NIR-II structured-illumination light-sheet microscopy. Proc. Natl Acad. Sci. USA 118, e2023888118 (2021).

20. Zhao, R. et al. A review of light-field imaging in biomedical sciences. Med-X 3, 25 (2025).

21. Zhao, R., Park, J. & Gao, L. Snapshot 3D at the speed frontier: redefining light-field microscopy for neuroimaging. PhotoniX 7, 46 (2026).

22. Gao, L. & Wang, L. V. A review of snapshot multidimensional optical imaging: measuring photon tags in parallel. Phys. Rep. 616, 1–37 (2016).

23. Colbert, A. E. et al. Enhanced infrared photodiodes based on PbS/PbClx core/shell nanocrystals. ACS Appl. Mater. Interfaces 13, 58916–58926 (2021).

24. Chen, H., Asif, M. S., Sankaranarayanan, A. C. & Veeraraghavan, A. FPA-CS: focal plane array-based compressive imaging in short-wave infrared. In Proc. IEEE Conference on Computer Vision and Pattern Recognition 2358–2366 (IEEE, 2015).

25. Antaris, A. L. et al. A high quantum yield molecule–protein complex fluorophore for near-infrared II imaging. Nat. Commun. 8, 15269 (2017).

26. Zhong, F., et al. Fast in vivo deep-tissue 3D imaging with selective-illumination NIR-II light-field microscopy and aberration-corrected implicit neural representation. Laser Photonics Rev. 10.1002/lpor.71525 (2026).

27. Wang, Z. et al. Kilohertz volumetric imaging of in vivo dynamics using squeezed light field microscopy. Nat. Methods 22, 2194–2204 (2025).

28. Fellgett, P. B. Conclusions on multiplex methods. J. Phys. Colloq. 28, C2-165–C2-171 (1967).

29. Gao, B. et al. NIR-II light field microscopy for through-skull hemodynamic volumetric imaging in awake mice. Laser Photonics Rev. 19, 2401685 (2025).

30. Cosco, E. D. et al. Bright chromenylium polymethine dyes enable fast, four-color in vivo imaging with shortwave infrared detection. J. Am. Chem. Soc. 143, 6836–6846 (2021).

31. Ng, R. Fourier slice photography. ACM Trans. Graph. 24, 735–744 (2005).

32. Harrison, M. R. M. et al. Late developing cardiac lymphatic vasculature supports adult zebrafish heart function and regeneration. eLife 8, e42762 (2019).

33. Harrison, M. R. M. et al. Chemokine-guided angiogenesis directs coronary vasculature formation in zebrafish. Dev. Cell 33, 442–454 (2015).

34. González-Rosa, J. M. Zebrafish models of cardiac disease: from fortuitous mutants to precision medicine. Circ. Res. 130, 1803–1826 (2022).

35. Poon, K. L. & Brand, T. The zebrafish model system in cardiovascular research: a tiny fish with mighty prospects. Glob. Cardiol. Sci. Pract. 2013, 9–28 (2013).

36. Lee, J. et al. 4-Dimensional light-sheet microscopy to elucidate shear stress modulation of cardiac trabeculation. J. Clin. Invest. 126, 1679–1690 (2016).

37. Wang, Z. et al. Real-time volumetric reconstruction of biological dynamics with light-field microscopy and deep learning. Nat. Methods 18, 551–556 (2021).

38. Bensimon-Brito, A. et al. Integration of multiple imaging platforms to uncover cardiovascular defects in adult zebrafish. Cardiovasc. Res. 118, 2665–2687 (2022).

39. Martin, R. T. & Bartman, T. Analysis of heart valve development in larval zebrafish. Dev. Dyn. 238, 1796–1802 (2009).

40. Berdougo, E., Coleman, H., Lee, D. H., Stainier, D. Y. R. & Yelon, D. Mutation of weak atrium/atrial myosin heavy chain disrupts atrial function and influences ventricular morphogenesis in zebrafish. Development 130, 6121–6129 (2003).

41. Kalogirou, S. et al. Intracardiac flow dynamics regulate atrioventricular valve morphogenesis. Cardiovasc. Res. 104, 49–60 (2014).

42. Wang, J. et al. A continuum of atrial peristalsis initiates the bicuspid to quadricuspid valve transition. Preprint at bioRxiv 10.64898/2026.05.04.722768 (2026).

43. Stacker, S. A. et al. Lymphangiogenesis and lymphatic vessel remodelling in cancer. Nat. Rev. Cancer 14, 159–172 (2014).

44. Kwon, S., Agollah, G. D., Wu, G., Chan, W. & Sevick-Muraca, E. M. Direct visualization of changes of lymphatic function and drainage pathways in lymph node metastasis of B16F10 melanoma using near-infrared fluorescence imaging. Biomed. Opt. Express 4, 967–977 (2013).

45. Honkura, N. et al. Intravital imaging-based analysis tools for vessel identification and assessment of concurrent dynamic vascular events. Nat. Commun. 9, 2746 (2018).

46. Dietzel, S. et al. Label-free determination of hemodynamic parameters in the microcirculation with third harmonic generation microscopy. PLoS ONE 9, e99615 (2014).

47. Pal, S. et al. Rhythmic contractions of lymph vessels and lymph flow are disrupted in hypertensive rats. Hypertension 82, 72–83 (2025).

48. Pal, S., Henry, D., Rhee, S. W. & Stolarz, A. J. Hypertension induces contractile dysfunction in rat mesenteric lymph vessels. FASEB J. 36 (Suppl. 1), abstract 0R509 (2022).

49. Wang, Z. et al. Methods, trade-offs and opportunities in high-speed optical microscopy for neural voltage imaging. Nat. Photon. 10.1038/s41566-026-01965-5 (2026).

50. Xiao, S., Tseng, H., Gritton, H., Han, X. & Mertz, J. Video-rate volumetric neuronal imaging using 3D targeted illumination. Sci. Rep. 8, 7921 (2018).

51. Xiao, S. et al. Large-scale deep tissue voltage imaging with targeted-illumination confocal microscopy. Nat. Methods 21, 1094–1102 (2024).

52. Mangeat, T. et al. Super-resolved live-cell imaging using random illumination microscopy. Cell Rep. Methods 1, 100009 (2021).

53. Mazzella, L. et al. Extended-depth of field random illumination microscopy, EDF-RIM, provides super-resolved projective imaging. Light Sci. Appl. 13, 285 (2024).

54. Briers, J. D. & Webster, S. Laser speckle contrast analysis (LASCA): a nonscanning, full-field technique for monitoring capillary blood flow. J. Biomed. Opt. 1, 174–179 (1996).

55. Ou, Z. et al. Achieving optical transparency in live animals with absorbing molecules. Science 385, eadm6869 (2024).

56. Feautrier, P., et al. C-RED 2 InGaAs 640x512 600-fps infrared camera for low order wavefront sensing. Proc. SPIE 10703, 107031V (2018).

57. Li, X. et al. Reinforcing neuron extraction and spike inference in calcium imaging using deep self-supervised denoising. Nat. Methods 18, 1395–1400 (2021).

58. Li, X. et al. Real-time denoising enables high-sensitivity fluorescence time-lapse imaging beyond the shot-noise limit. Nat. Biotechnol. 41, 282–292 (2023).

